# Interpretable spherical geometry of single-cell state transitions from dominant principal components

**DOI:** 10.64898/2026.09.11.751061

**Authors:** Long Yuan, Xuyang Li, My Le, Stephanie C. Hicks, Atul Deshpande, Janis M. Taube, Alexander S. Szalay

**Author notes:** **Corresponding authors:** To whom correspondence may be addressed. (L.Y.), (J.M.T), or (A.S.S).

## Abstract

Single-cell RNA-seq atlases are commonly explored with nonlinear embeddings that preserve neighborhoods but provide limited coordinate-level interpretation. We asked whether projecting the dominant principal components (PCs) of single-cell gene expression onto a unit sphere would yield an interpretable coordinate system. SPHERE-PCA *L*_2_-normalizes the first three PC coordinates, aligns a biologically defined root to the north pole, and represents each cell by three coordinates: root-aligned geodesic distance (*θ*), angular position (*ϕ*), and pre-projection radial magnitude (*r*). Across developmental and disease-associated datasets, this representation reveals structured spherical geometry, ranging from near-great-circle trajectories to multi-arc manifolds. In developmental atlases, root-aligned geodesic distance increases as CytoTRACE-inferred stemness decreases, while gene-coordinate analyses separate programs associated with angular position from those associated with radial magnitude. Fixed-loading perturbations decompose each gene’s effect on cell position into progression, branch- or state-position, and radial activity components. SPHERE-PCA therefore provides a deterministic, loading-preserving coordinate framework for interpreting dominant transcriptomic variance and establishing a transparent geometric coordinate framework for perturbation analysis and virtual-cell models.

**Significance Statement:** Single-cell atlases are often interpreted through nonlinear maps whose axes are visually powerful but difficult to explain. This study shows that projecting the first three principal components onto a unit sphere yields an interpretable coordinate system for single-cell data. Across developmental atlases, the resulting coordinates expose distance from a biological root, angular branch or state position, and pre-projection radial magnitude, which capture progression-associated, state-associated, and activity-associated variation. Because the method preserves principal component loadings, gene-associated displacements can be decomposed along the same coordinates. SPHERE-PCA provides a transparent geometric layer on classical dimension reduction and a coordinate scaffold for future models of cell-state change such as virtual-cell.

## Introduction

Single-cell atlases are often explored through low-dimensional maps. Methods such as UMAP and t-SNE have become central tools for visualizing neighborhoods, separating cell states, and communicating complex transcriptomic structure [1, 2]. Trajectory and graph-abstraction methods provide complementary frameworks for ordering cells along inferred transitions [3–5]. Yet the axes of many widely used embeddings do not correspond directly to biological quantities, and recent work has emphasized that embedding metrics can be misleading when global and local structure are not distinguished [6, 7]. This creates a practical need for representations whose axes, distances, and gene associations can be described directly.

Principal component analysis (PCA) remains one of the most transparent ways to summarize transcriptomic variance, but PCA plots of continuous biological gradients often curve into arcs, horseshoes, or oscillatory patterns. Such effects have long been studied in ordination, multidimensional scaling, microbial community analysis, population genetics, and single-cell RNA-seq [8–13]. These shapes are often treated as artifacts to diagnose or avoid. We asked instead whether, in datasets where dominant PCA curvature is stable and biologically anchored, projecting the first three principal components onto a sphere could transform the curved geometry of a PCA plot into a coordinate system with independently interpretable axes.

Geometric embeddings have precedent in single-cell analysis. scPhere uses deep generative modeling to learn spherical and hyperbolic latent spaces [14], and related non-Euclidean approaches have been used to represent hierarchical relationships and cross-dataset correspondence [15, 16]. SPHERE-PCA differs in design. It trains no deep model, introduces no nonlinear optimization objective, and preserves the PCA loading matrix. The method uses dominant PCA coordinates, projects each cell vector to the unit sphere, and retains the pre-projection magnitude as a separate scalar. After choosing a biological root or reference state, root-aligned geodesic distance, angular position around the root, and pre-projection radial magnitude can be analyzed as distinct components of cell-state variation.

Here we apply SPHERE-PCA to a range of datasets, spanning ordered differentiation, disease-associated heterogeneity, and targeted spatial transcriptomics, to test whether dominant PC geometry is interpretable across contexts. We test whether this geometry is robust across datasets, whether *θ* tracks transcriptomic stemness scores derived from an independent algorithm, and whether gene programs separate into angular and radial components. Finally, because SPHERE-PCA remains tied to PCA loadings, we introduce a fixed-loading perturbation analysis that decomposes each gene’s contribution into progression-associated movement, branch/state-position displacement, and pre-projection radial magnitude change. Together, these results establish SPHERE-PCA as a deterministic, loading-preserving geometric framework for interpreting dominant transcriptomic variance, complementing nonlinear visualization and specialized trajectory inference methods.

## Results

### SPHERE-PCA maps dominant transcriptomic variation onto interpretable spherical coordinates

SPHERE-PCA takes as input a cell-by-gene expression matrix or precomputed PCA scores, uses the first three principal components as a 3D coordinate, and *L*_2_-normalizes each coordinate to unit length (Fig. 1A). For cell *i* with PC coordinate *x_i_* = (*PC*1*_i_, PC*2*_i_, PC*3*_i_*), the spherical coordinate is *x̂_i_* = *x_i_/*∥*x_i_*∥. The projection preserves directional information in PC1-3 space while discarding pre-projection radial magnitude from the sphere; this magnitude is retained as an auxiliary scalar. Longitude-latitude plots are then equirectangular visualizations of the sphere, not separate embeddings.

**Figure 1:**
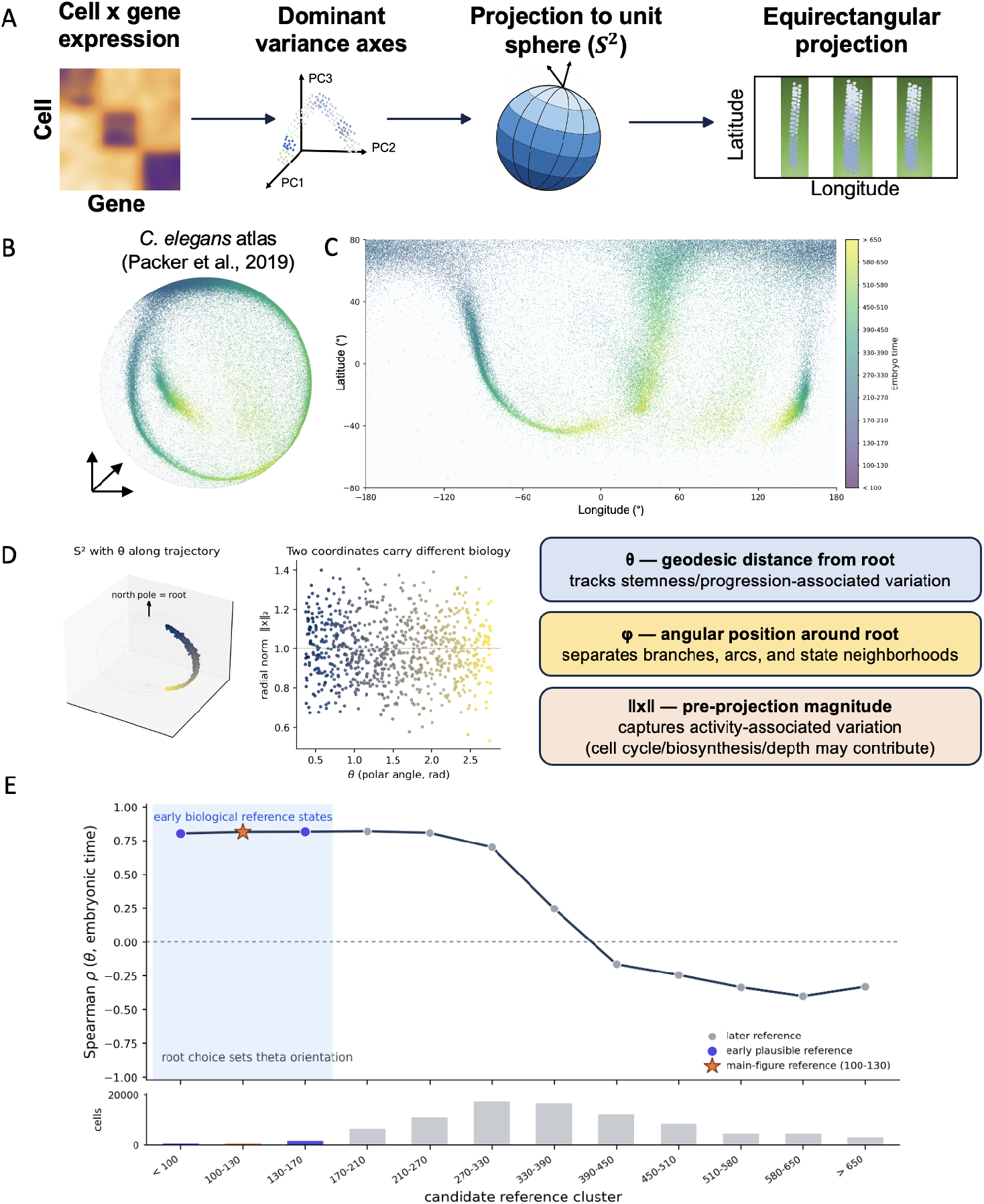
SPHERE-PCA maps dominant single-cell transcriptional variation onto a sphere and separates root distance, angular position, and radial magnitude. (A) Conceptual workflow. A cell-by-gene expression matrix is projected onto dominant PCA axes, the first three PC coordinates are *L*_2_-normalized onto the unit sphere *S*^2^, and spherical coordinates can be visualized by equirectangular projection after optional rotation. (B) *C. elegans* embryogenesis atlas represented on the unit sphere. Points are cells colored by embryonic time bin. (C) Equirectangular projection of the same root-aligned embedding. Longitude and latitude display stripe-like arcs associated with embryonic time; annotated stripe bands are illustrative visual guides. (D) Coordinate interpretation schematic. After choosing a biological root, *θ* denotes root-aligned geodesic distance, *ϕ* denotes angular position around the root, and ∥*x*∥ denotes pre-projection PC1-3 magnitude retained as an auxiliary scalar. (E) Root/reference sensitivity in the *C. elegans* atlas. For each candidate time bin, *θ* was recalculated as geodesic distance from that reference and correlated with ordered embryonic time by Spearman correlation. Early roots preserve the expected orientation, whereas later roots can weaken or flip the association.

Applied to the *C. elegans* embryogenesis atlas, SPHERE-PCA revealed arc-like, time-stratified organization when cells were colored by embryonic time bins (Fig. 1B and C). The reference cluster was the earliest embryonic time bin (100-130 min), consistent with the root convention used for related spherical developmental analyses [14]. After aligning this reference to the north pole and applying a fixed Euler rotation for visualization, the 3D sphere and its equirectangular projection revealed repeated time-associated bands (Fig. 1C).

The coordinate interpretation is root dependent. We define *θ* as root-aligned geodesic distance from the chosen biological reference, *ϕ* as angular position around the root axis, and ∥*x*∥ as pre-projection PC1-3 radial magnitude (Fig. 1D). Thus *θ* is not pseudotime by definition; it is a distance coordinate that can be tested against developmental time or known state order. Similarly, *ϕ* may separate branch or state neighborhoods but does not by itself assign lineage fate. The pre-projection radial magnitude is removed by *L*_2_-normalization but is retained as an auxiliary scalar; it may reflect activity-associated variation such as cell-cycle state, biosynthetic output, or broad transcriptional activation, and may also absorb technical variation related to sequencing depth.

Root sensitivity analysis quantified how the *θ*-time relationship depends on the chosen reference state (Fig. 1E). For each candidate embryonic time bin, the root centroid was aligned to the north pole and *θ* was correlated with ordered embryonic time. Early reference states showed positive correlations with embryonic time, including the chosen 100-130 bin (*ρ* = 0.814); later references weakened or reversed the association, with bins after 390 min showing negative correlations. This result shows that any biological interpretation of *θ* requires explicit reporting of the chosen reference state, since the direction of the developmental association depends on which reference is used.

### Dominant PC geometry forms robust spherical structures across datasets

We next asked whether spherical structure was specific to one dataset or to a single display orientation. Across datasets, the dominant PCA geometry ranged from a single near-global arc to multi-arc organization with multiple visible branches (Fig. 2). A single great-circle fit summarized the mouse embryonic stem cell (ESC) dataset well (*R*^2^ = 0.936) but was only a baseline for the more complex *C. elegans* atlas (*R*^2^ = 0.790), where multiple arcs remained visible (Fig. 2A) [17, 18].

**Figure 2:**
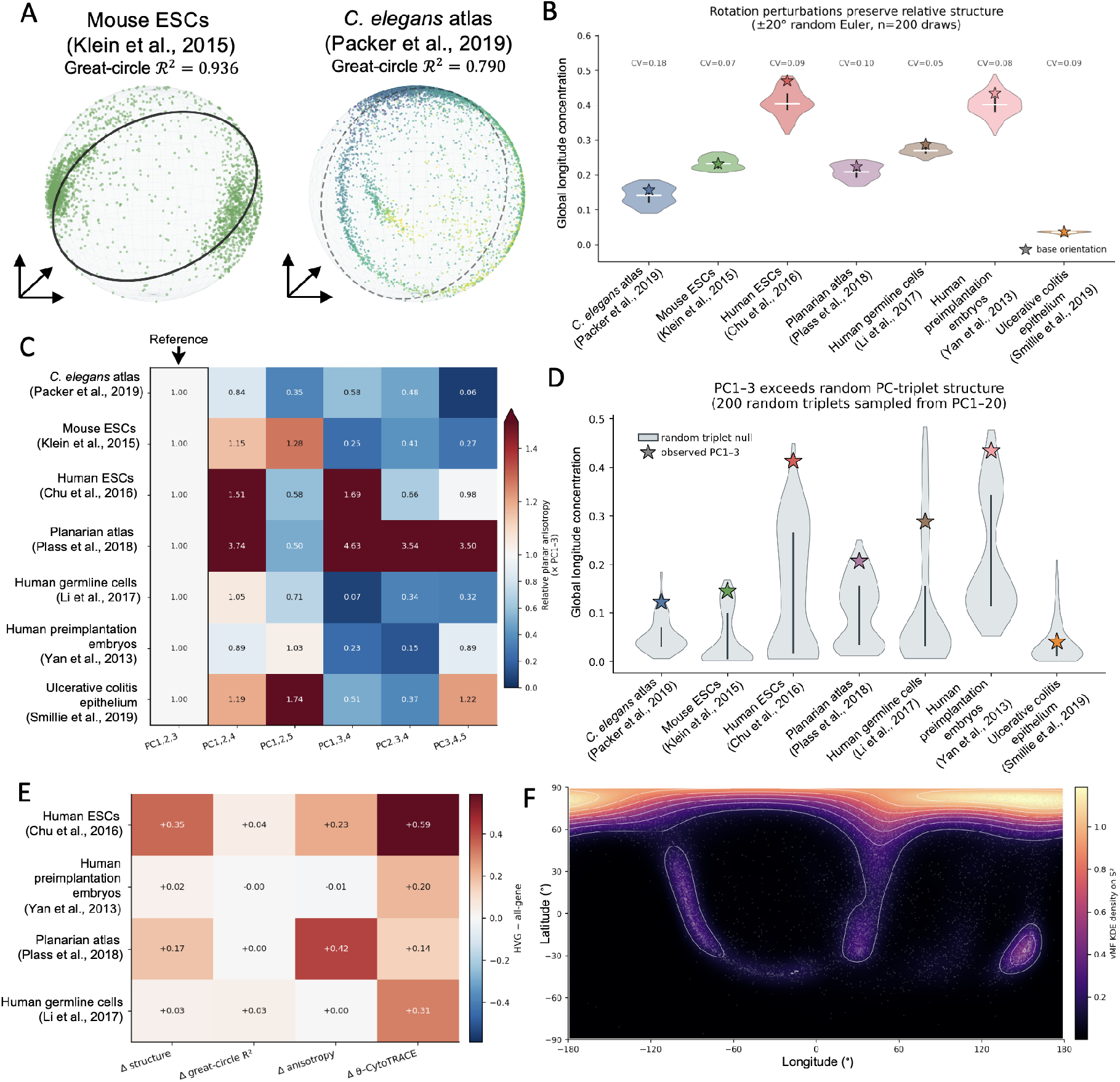
Dominant PC geometry forms robust spherical structure ranging from global trajectories to multi-arc manifolds. (A) Great-circle baseline fits for mouse ESCs and the *C. elegans* embryogenesis atlas. Cells are plotted on *S*^2^; the overlaid great circle summarizes near-global geometry in mouse ESCs but is only a baseline for the multi-arc *C. elegans* atlas. (B) Rotation perturbation analysis. For each dataset, global longitude concentration was recomputed after 200 random 20*^◦^* Euler perturbations. Violin distributions show perturbation scores, with the base orientation indicated. (C) PC-axis specificity. Structure scores for alternative PC triplets are normalized to the PC1-3 score. Triplets retaining early dominant axes frequently preserve more structure than later-PC combinations, although alternatives can exceed PC1-3 in some datasets. (D) Random PC-triplet null analysis. Observed PC1-3 structure is compared with 200 random triplets sampled from PC1-PC20. This is a geometric null, not a biological null. (E) Feature-selection robustness. HVG-based PCA and all-gene PCA are compared for structure, great-circle fit, anisotropy, and stemness association. (F) Spherical density visualization of the *C. elegans* embedding. von Mises–Fisher KDE on the equirectangular projection highlights multiple local density ridges, supporting multi-arc structure without assigning a definitive lineage count.

Small rotations did not destroy the observed structure. For seven datasets, we applied 200 random Euler perturbations within 20*^◦^* and recomputed global longitude concentration, a scalar summary of non-uniform angular structure (Fig. 2B). Base orientations lay within the perturbation envelopes, indicating that the observed angular structure did not depend on a fine-tuned viewing orientation. This perturbation analysis tests robustness to small coordinate or viewing changes, not whether all orientations are biologically equivalent.

The structure was also specific to dominant PC axes rather than arbitrary higher-index PCs. PC-triplet comparisons showed that triplets containing low-index axes often showed stronger structure than combinations of higher-index PCs, although some datasets had alternative low-index triplets that exceeded PC1-3 in structure score (Fig. 2C). This is expected for complex data: PC1-3 is a transparent default, not a theorem of optimality. Against a random PC-triplet null sampled from PC1-20, observed PC1-3 scores were above the null mean in all seven datasets (Fig. 2D). The signal was strongest in hESCs and preimplantation embryos and weakest in ulcerative colitis epithelium as expected, given that developmental systems have clearer progression hierarchies than disease or inflammatory contexts, where multiple cell states coexist without a necessary defined ordering[19–21].

Feature selection modulated quantitative metrics but did not alone account for the observed geometry (Fig. 2E). In hESCs, planaria, human germ cells, and preimplantation embryos, Seurat-style highly variable gene (HVG) selection modulated structure, great-circle fit, planar anisotropy, and stemness correlations compared with all-gene PCA [22]. Finally, von Mises-Fisher kernel density estimation on the *C. elegans* sphere highlighted multiple local density ridges (Fig. 2F). We interpret these ridges as evidence for multi-arc spherical organization, not as a definitive enumration of biological lineages.

### Root-aligned spherical distance tracks transcriptomic stemness and separates angular and radial gene programs

To test biological interpretation directly, we aligned SPHERE-PCA embeddings to pre-specified stem-like or early reference states and compared root-aligned geodesic distance with CytoTRACE-inferred differentiation state, an algorithmically distinct transcriptional-diversity-based measure [23]. The roots were hESC for human ESC differentiation, x1/neoblast for planaria, 19W for human germ cells, and zygote for human preimplantation embryos. These roots were chosen a priori from biological annotations as stem-like or earliest available states and were associated with high CytoTRACE-inferred developmental potential, consistent with their stem-like annotation.

We first used known planarian lineages as a labeled test of angular organization. Six curated lineage paths rooted at neoblast 1 occupied compact angular sectors on the sphere (Fig. 3A; Supplementary Fig. S1B)[24]. Along these known paths, Spearman correlations between published lineage order and *θ* were positive for all tested lineages, ranging from 0.353 for gut to 0.675 for epidermal (Fig. 3B). Label-shuffling permutation tests found stronger within-lineage angular compactness than expected by chance for all six lineages (*P <* 0.001, 1000 permutations; Fig. 3C). Centroid-direction heatmaps comparing angular distances in 3D SPHERE-PCA space with angular distances in PC1–50 space showed that known lineage groups are more visually separable in the 3D spherical view, though this does not imply that SPHRER-PCA outperforms higher-dimensional PCA representations across all settings (Supplementary Fig. S1A). Together, these analyses show that SPHERE-PCA angular coordinates are consistent with known planarian lineage annotations.

**Figure 3:**
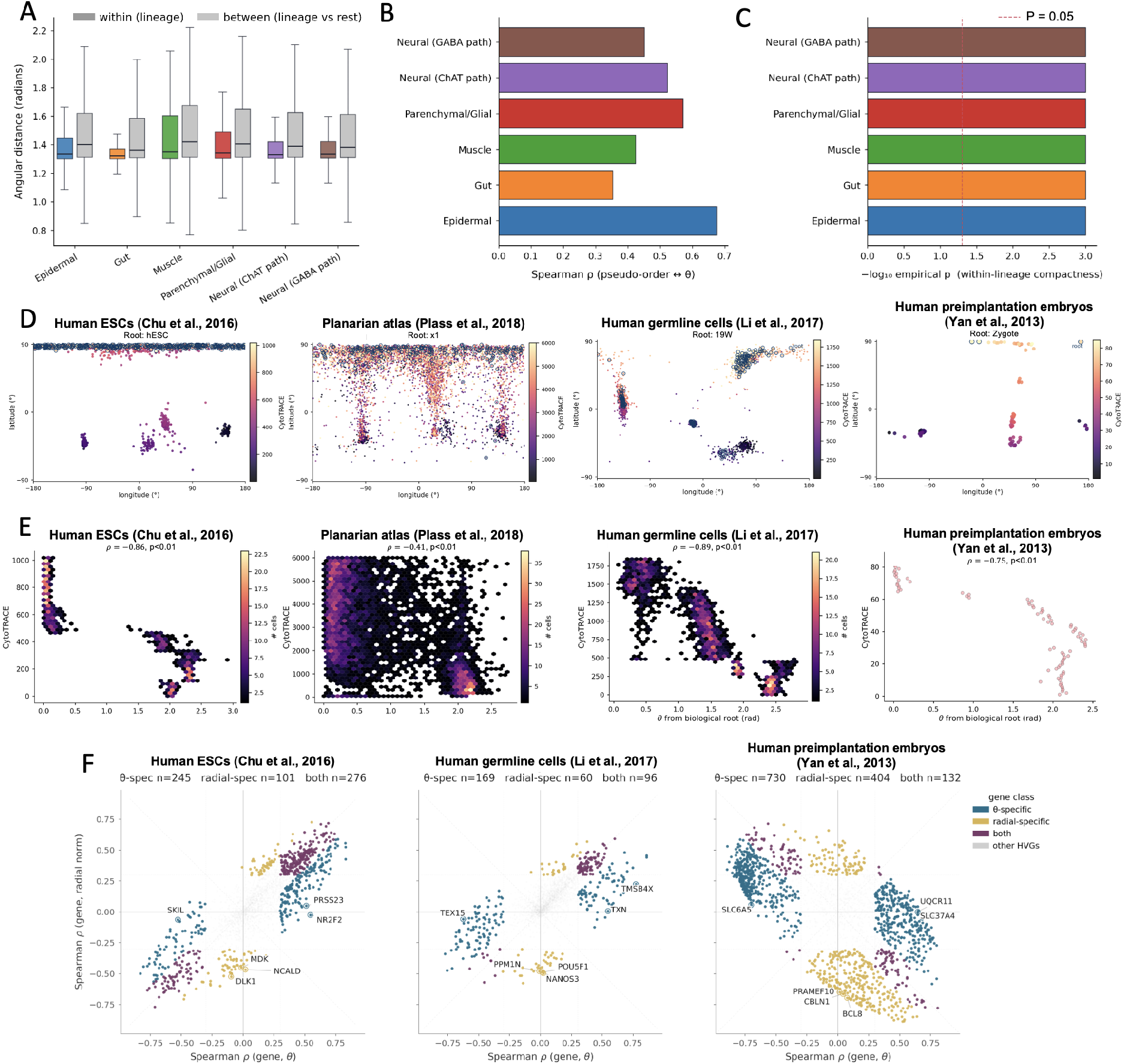
Spherical distance from a biologically defined stem-like root tracks loss of transcriptomic stemness. (A) Known planarian lineages occupy angularly compact sectors on the SPHERE-PCA embedding. Boxplots compare within-lineage angular distances with between-lineage-versus-rest angular distances for curated lineage groups. (B) Spearman correlation between curated planarian lineage order and root-aligned geodesic distance from the neoblast reference. Positive correlations indicate increasing *θ* along known lineage paths. (C) Label-shuffling permutation tests for lineage angular compactness. Bars show − log_10_ empirical permutation *P*; the dashed line marks *P* = 0.05. (D) Root-aligned SPHERE-PCA embeddings colored by raw CytoTRACE score for human ESCs, planaria, human germ cells, and human preimplantation embryos. Roots were biologically pre-specified. (E) Quantitative relationship between *θ* and CytoTRACE. Because *θ* increases away from high-stemness roots, negative Spearman correlations indicate loss of transcriptomic stemness with increasing root-aligned spherical distance. (F) Gene-coordinate specificity. Each point is an HVG plotted by its Spearman correlation with *θ* and with pre-projection radial norm. Colors denote *θ*-specific, radial-specific, both-coordinate, and other genes. Gene-coordinate associations are correlative.

Across four CytoTRACE datasets, high-stemness cells were concentrated near the selected root in root-aligned equirectangular views (Fig. 3D). Quantitatively, CytoTRACE-inferred developmental potential was negatively correlated with *θ* in all four datasets (Fig. 3E): hESCs (*ρ* = −0.858, *n* = 1018), planaria (*ρ* = −0.411, *n* = 6000), human germ cells (*ρ* = −0.892, *n* = 1844), and preimplantation embryos (*ρ* = −0.751, *n* = 85). The negative sign is expected because geodesic distance increases away from a high-stemness root. These results support the interpretation of root-aligned geodesic distance as progression-associated in these datasets, while treating CytoTRACE as an complementary algorithmic measure rather than an independent ground truth.

Gene-coordinate analysis further separated genes into spherical- and radial-coordinate classes (Fig. 3F). For hESCs, human germ cells, and preimplantation embryos, each HVG was correlated with *θ* and with pre-projection radial norm. Genes were classified using *|ρ| ≥* 0.30, coordinate-specificity difference *≥* 0.15, and false discovery rate (FDR) *<* 0.05. The resulting classes included *θ*-specific genes, radial-specific genes, and genes associated with both coordinates. Across the three datasets, hundreds of HVGs fell into coordinate-specific classes, with class sizes dvarying by dataset and root. These correlations are descriptive, but they indicate that *L*_2_ projection can preserve directional programs on the sphere while separating magnitude-associated variation into the pre-projection radial scalar. Enrichment analyses and overlap with the canonical Tirosh et al. cell-cycle marker list further supported the activity-associated interpretation of the radial coordinate, with the strongest overlap in the hESC both-coordinate class (Supplementary Fig. S2) [25].

### Fixed-loading gene perturbations decompose progression, branch/state position, and radial activity components

Because SPHERE-PCA is built directly from PCA loadings, individual genes can be analyzed in the same coordinate system. For each HVG *g*, we performed a fixed-loading perturbation in PC space,

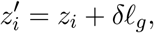

where *z_i_* is the cell’s PC1-3 coordinate, *ℓ_g_* is the gene’s loading vector on PC1-3, and *δ* = 1.0 corresponds to a +1 SD perturbation in standardzed expression unites (see Methods). We then projected perturbed coordinates back to the root-aligned sphere and summarized each gene by median Δ*θ*, median *|*Δ*ϕ|*, and median Δ log_2_ *r*. These quantities capture computational displacements in fixed PCA-loading space and should not be interpreted as predictions of experimental perturbation outcomes.

In the hESC dataset, selected genes illustrated each of the three coordinate components (Fig. 4A–C). For each gene, we applied a fixed *δ* = 1.0 perturbation along its PC1–3 loading vector and summarized the resulting displacement by progression-associated shift

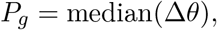

branch/state-position shift

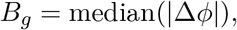

and radial shift

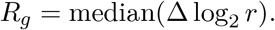

*COL5A2* showed positive progression-associated displacement (*P_g_* = 0.106*^◦^*), *PTPRZ1* showed counter-progression displacement (*P_g_* = −0.111*^◦^*), and *FZD5* showed strong branch/state-position displacement (*B_g_* = 0.713*^◦^*). Arrows in Fig. 4A are visually amplified 30-fold and should not be interpreted as biological effect sizes.

**Figure 4:**
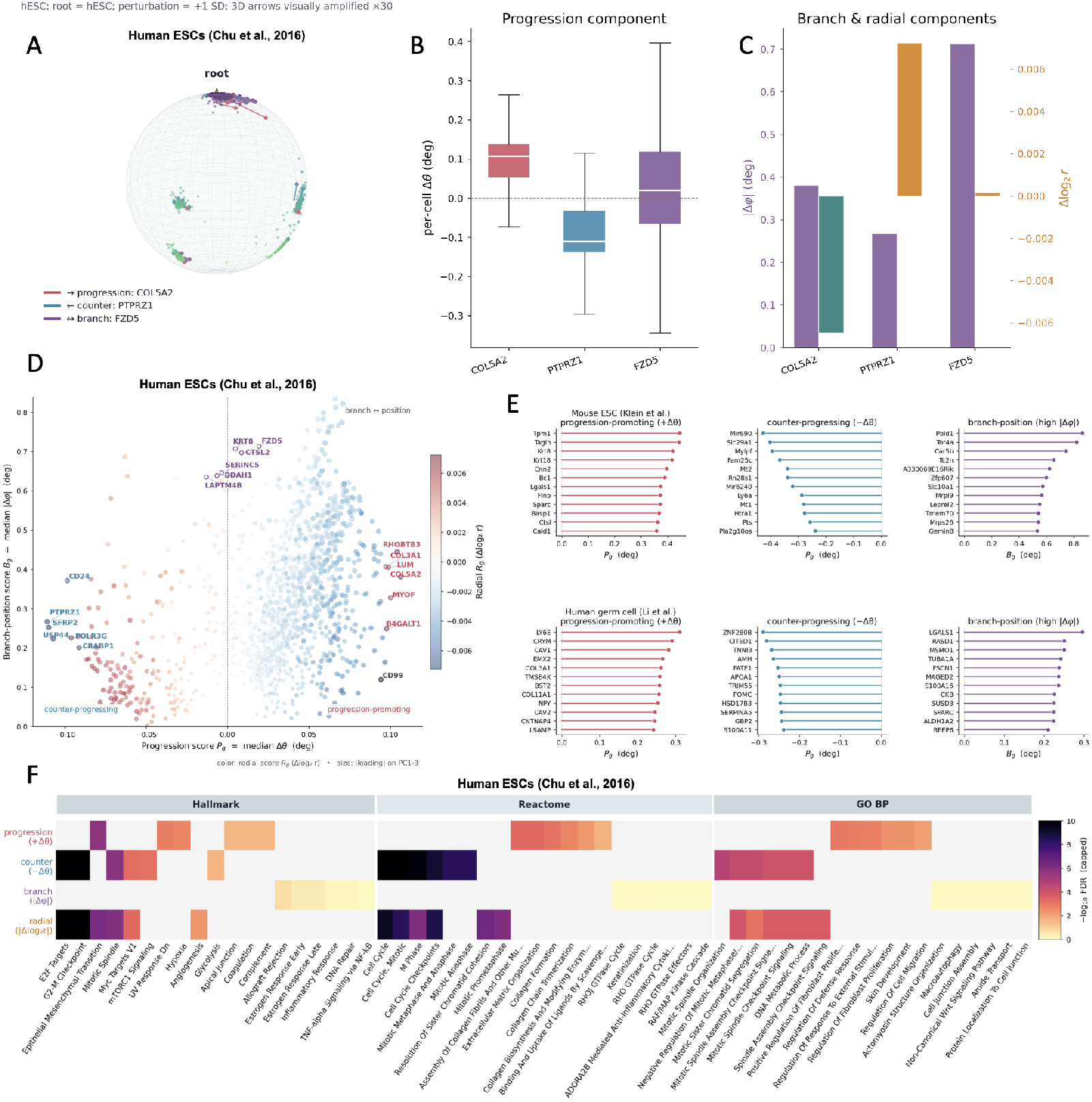
Gene perturbations decompose progression, branch/state position, and activity on the sphere. (A) Example fixed-loading gene perturbations on the hESC SPHERE-PCA embedding. For each selected gene, a +1 SD perturbation was applied in the standardized gene-expression space, equivalently *δ* = 1 after scaling, and propagated through the gene’s fixed PC1–3 loading vector before re-projecting cells onto the root-aligned sphere. Arrows are visually amplified for display only; all statistics are computed from the unamplified *δ* = 1 perturbation. (B) Per-cell progression component Δ*θ* for selected example genes. Positive values indicate movement away from the hESC root, whereas negative values indicate movement toward the root. (C) Branch/state-position and radial components for selected examples. Purple bars show the branch/state-position component *|*Δ*ϕ|*, which captures lateral displacement around the sphere. Orange/teal bars show the radial component Δ log_2_ *r*, which captures changes in pre-projection radial magnitude; positive and negative values indicate increased and decreased radial magnitude, respectively. (D) hESC gene perturbation map. Each gene is plotted by progression score *P_g_* and branch/state-position score *B_g_*, with color indicating radial score *R_g_* and point size indicating PC1-3 loading magnitude. (E) Coordinate-specific gene rankings in mouse ESCs and human germline cells for progression, counter-progression, and branch/state-position components. (F) Pathway enrichment of coordinate-specific gene classes using Hallmark, Reactome, and Gene Ontology Biological Process gene sets. Color shows − log_10_ FDR capped at 10.

Mapping all analyzed hESC HVGs showed that genes separated into coordinate-specific regions (Fig. 4D). Progression-associated genes with positive median Δ*θ* included *COL5A2*, *RHOBTB3*, *MYOF*, *LUM*, *B4GALT1*, and *COL3A1*, consistent with extracellular-matrix-associated differentiation or remodeling programs [26]. Genes with negative median Δ*θ* (counter-progressing toward the root) included *PTPRZ1*, *SFRP2*, *USP44*, *CRABP1*, *POLR3G*, and *CD24*. Among these, *POLR3G* and *CD24* have been linked to human pluripotent stem-cell maintenance [27, 28]. Genes with high *|*Δ*ϕ|*, including *FZD5*, *KRT8*, *CTSL2*, *SERINC5*, *DDAH1*, and *LAPTM4B*, pointed to branch- or state-position-associated programs, including Wnt/FZD and epithelial or membrane-associated signals [29]. Together, these enrichment patterns support the biological coherence of the SPHERE-PCA coordinate decomposition.

The same scoring procedure produced coordinate-specific rankings in mouse ESCs and human germ cells (Fig. 4E). The top-ranked genes were dataset-specific, as expected across species, but the coordinate decomposition itself transferred readily across datasets. In hESCs, pathway enrichment of coordinate classes against the HVG background showed distinct programs (Fig. 4F). Progression-associated genes were enriched for extracellular matrix, collagen, epithelial–mesenchymal transition, and related matrix-remodeling terms, whereas counter-progressing and radial classes were enriched for E2F, G2/M, mitotic, and broader cell-cycle programs. Branch/state-position genes showed weaker but biologically interpretable enrichment for lineage- or state-context terms, including signaling, response, localization, and junction-associated processes. Together, these patterns support the biological coherence of SPHERE-PCA coordinate classes and nominate candidate programs for future functional validation.

### Context-dependent structure and benchmarking beyond developmental atlases

Supplementary analyses tested where SPHERE-PCA is informative and where interpretation requires caution. Non-developmental scRNA-seq and targeted spatial-transcriptomic datasets served as stress tests for method generality rather than as primary evidence for developmental progression (Supplementary Fig. S3). A breast cancer scRNA-seq atlas produced cell-type-associated regions on the sphere, suggesting that dominant PC geometry can organize broad state variation outside embryogenesis (Supplementary Fig. S3A) [30]. We next analyzed data from Xenium, an imaging-based spatial transcriptomic platform that measures targeted gene panels rather than whole-transcriptome profiles, including breast cancer and lung cancer panels before and after TRACER, a mutual-information-based transcript-assignment and segmentation-refinement method (Supplementary Fig. S3B–F) [31–33]. These targeted panels, each containing approximately 300 genes, nevertheless produced visible spherical structure. After TRACER refinement, the spherical embeddings showed sharper stripe-like organization, consistent with improved transcript-to-cell assignment or reduced segmentation-associated noise. Because these datasets are targeted spatial panels rather than whole-transcriptome developmental atlases, their geometry reflects spatial distribution and cell-type composition, and is also influenced by the targeted gene panel.

Finally, a *C. elegans* benchmark compared PCA (PC1–3), SPHERE-PCA, 3D UMAP, 3D t-SNE, and a precomputed 3D scPhere latent embedding against a PC1–50 reference (Supplementary Fig. S4). UMAP and t-SNE showed higher trustworthiness, continuity, and k-nearest-neighbor Jaccard overlap, consistent with their local-neighborhood objectives. SPHERE-PCA preserved a structured global developmental gradient, with *|ρ|* = 0.782 between anchor distance and embryonic time, compared with 0.738 for PCA, 0.820 for UMAP, 0.502 for t-SNE, and 0.857 for scPhere. Thus, the benchmark supports complementary behavior: SPHERE-PCA provides an interpretable global coordinate while retaining measurable neighborhood structure, but it does not dominate nonlinear embeddings or probabilistic latent-variable models across all metrics.

## Discussion

The main conceptual result of this study is that dominant PCA curvature can be interpreted through spherical geometry when the coordinate system is rooted, tested, and reported explicitly. SPHERE-PCA does not remove the arc-like structure often seen in PCA plots. Instead, it maps the first three PC axes to *S*^2^, conveting the directional structure in PC space into a root-aligned geodesic spherical coordinate system and separating it from the pre-projection radial magnitude. In developmental systems, this produced a useful decomposition: root-aligned geodesic distance tracked progression-associated loss of transcriptomic stemness, angular position organized branch or state neighborhoods, and radial magnitude captured activity-associated variation.

Root choice is central. The sign and interpretation of *θ* depend on the selected reference, just as pseudotime depends on a root in trajectory inference. The *C. elegans* sensitivity analysis shows that early reference states maintain the expected orientation of *θ* relative to embryonic time, whereas later references can flip or weaken the association. This should be treated as an explicit modeling constraint rather than a source of hidden flexibility. A SPHERE-PCA analysis should state the root, justify it biologically, and test coordinate interpretation against external labels or scores whenever possible.

The method is intentionally constrained in scope relative to nonlinear and deep embeddings. UMAP, t-SNE, pseudotime algorithms, scPhere, and other geometric models solve different problems and can preserve local neighborhoods, branch topology, or learned latent manifolds more effectively in specific settings [5, 6, 14]. SPHERE-PCA is useful because it is deterministic, loading-preserving, and easy to audit. It should be interpreted as a geometric layer on dominant transcriptomic variance, not as a complete trajectory inference algorithm or a method for de novo lineage discovery.

The fixed-loading perturbation analysis extends this transparency to genes. Because each gene has an explicit loading vector, a gene perturbation in fixed PCA-loading space can be decomposed into movement along root-aligned geodesic distance, lateral movement around the sphere, and change in pre-projection radial magnitude. This analysis is not a substitute for perturb-seq or other experimental perturbation assays. It is a transparent linear analysis that characterizes whether a gene is associated with progression, counter-progression, branch- or state-position displacement, or radial activity-associated variation. Recent analyses have shown that simple linear baselines remain competitive with learned perturbation models [34], and SPHERE-PCA provides one such interpretable linear decomposition.

More broadly, SPHERE-PCA may provide a coordinate scaffold for evaluating whether virtual-cell or flow-field models learn biologically meaningful directions of cell-state movement. In this setting, learned perturbation vectors could be decomposed into progression-associated geodesic displacement, branch- or state-position angular displacement, and radial activity-associated displacement. The present study establishes this geometric coordinate system and demonstrates that fixed-loading gene displacements can be summarized within it, providing a transparent baseline for future learned perturbation models.

Several limitations are inherent to the approach. First, PC1-3 cannot contain all biologically relevant variation, and alternative early PC triplets may be more structured in some datasets. Second, radial magnitude can include biological activity (including cell-cycle state and biosynthetic output),technical factors such as sequencing depth, and compositional variation. Third, disease-associated and targeted spatial panels may show spherical organization without a developmental interpretation. Fourth, benchmark results show trade-offs, not universal superiority. Finally, gene-coordinate associations and fixed-loading perturbations are correlative and computational; causal interpretation requires experimental validation.

## Materials and Methods

### Datasets and data availability

Analyses used published single-cell and spatial transcriptomic datasets spanning developmental, stem/progenitor, inflammatory, cancer, and targeted spatial-transcriptomic contexts. These included *C. elegans* embryogenesis [18], mouse ESC differentiation [17], human ESC differentiation [19], planarian regeneration atlas [24], human germline cells [35], human preimplantation embryos [20], ulcerative colitis epithelium [21], a breast cancer scRNA-seq atlas [30], and Xenium breast cancer and lung cancer datasets analyzed before and after TRACER transcript-assignment refinement [31–33].

Processed *C. elegans* embryogenesis and ulcerative colitis epithelium data derived from scPhere [14] were downloaded from the Broad Institute Single Cell Portal, study SCP551: https://singlecell.broadinstitute.org/single_cell. The mouse ESC dataset is available from Gene Expression Omnibus under accession GSE65525. For the planarian atlas, we used the processed version available through the Planarian Schmidtea cell atlas portal, including the first 50 principal components: https://shiny.mdc-berlin.de/psca/. The breast cancer scRNA-seq atlas was downloaded from the Broad Institute Single Cell Portal, study SCP1039: https://singlecell.broadinstitute.org/single_cell/study/SCP1039. CytoTRACE-annotated versions of the human ESC differentiation, planarian, human germline, and human preimplantation embryo datasets were obtained from the CytoTRACE data portal: https://cytotrace.stanford.edu/. The multimodal Xenium lung cancer dataset, case TSU-20 section 1, was obtained from https://kero.hgc.jp/Ad-SpatialAnalysis_2024.html. The standard Xenium human breast cancer preview dataset was obtained from 10x Genomics: https://www.10xgenomics.com/products/xenium-in-situ/preview-dataset-human-breast.

### Preprocessing and PCA

Where raw expression matrices were available, cells were represented as AnnData objects and processed with Scanpy [36]. The canonical preprocessing pipeline consisted of: (1) total-count normalization to 10^4^ counts per cell; (2) log(1 + *x*) transformation; (3) highly variable gene (HVG) selection using the Seurat-style Scanpy implementation (scanpy.pp.highly_variable_genes(…, flavor=“seurat”)) with *n* = 2000 genes or min(2000*, n*_var_ − 1) when fewer genes were available; (4) scaling with values clipped at 10; and (5) PCA with zero_center=True and random seed 0 [22]. Unless otherwise specified, PCA was computed with 20 components and PC1–3 were used for SPHERE-PCA. Benchmarking and Xenium analyses used 50 computed PCs to support comparison with higher-dimensional reference spaces. For datasets distributed only as precomputed PCA coordinates, the provided PCs were used directly and the corresponding source is noted in the dataset description.

### SPHERE-PCA embedding and coordinates

For each cell, selected PC coordinates *x_i_ ∈* R^3^ were projected to the unit sphere by row-wise *L*_2_ normalization,

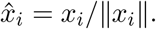

A root centroid was computed from the selected reference cells, normalized to unit length, and rotated to [0, 0, 1] using Rodrigues’ rotation formula. After root alignment, spherical coordinates were computed as *θ* = arccos(*z*) and *ϕ* = atan2(*y, x*), where *θ* is the polar angle from the root-aligned north pole and *ϕ* is the azimuthal angle. Equirectangular views used longitude = atan2(*y, x*) and latitude = arcsin(*z*). Fixed Euler rotations were used only for visualization unless explicitly tested; Fig. 1 used [−30, 0*, −*50]*^◦^*, and planaria supplemental visualizations used [80, 50, 50]*^◦^*.

### Structure, robustness, and null analyses

Global longitude concentration summarized non-uniform angular structure and was used as a relative structure score rather than a biological score. Longitude concentration was computed from the distribution of *ϕ* values after binning or circular-density estimation, with higher values indicating stronger angular non-uniformity. Great-circle fits estimated the best-fitting plane through the origin for spherical coordinates and returned *R*^2^ and residual angular distances from the fitted great circle. Spherical anisotropy was computed from the eigenvalues of the 3D scatter matrix of unit-sphere coordinates, with higher anisotropy indicating more directional concentration of points. Local density in *C. elegans* used von Mises–Fisher kernel density estimation with *κ* = 90, grid size 5000, 180 longitude bins, 90 latitude bins, and 7 contour levels. Rotation robustness used 200 random Euler perturbations within 20*^◦^*, a maximum of 8000 cells per dataset, and random seed 0. PC-triplet specificity compared specified triplets among PC1–PC5 or PC1–PC6 and normalized scores to PC1–3. Random null analyses sampled 200 triplets from PC1–PC20.

### Stemness, gene-coordinate, and lineage analyses

CytoTRACE values were obtained from the precomputed datasets described in the data availability section [23]. Root states were selected from biological annotations as stem-like or earliest available states: hESC for human ESC differentiation, x1/neoblast for planaria, 19W for human germ cells, and zygote for human preimplantation embryos. Spearman correlations between root-aligned polar angle *θ* and CytoTRACE were computed with scipy.stats.spearmanr [37]. Per-gene coordinate associations used log-normalized, non-scaled HVG expression and vectorized Spearman correlations with *θ* and pre-projection radial norm. Gene classes were defined using *|ρ| ≥* 0.30, coordinate-specificity difference *≥* 0.15, and Benjamini–Hochberg FDR *<* 0.05. Planarian lineage-sector statistics used six curated lineages rooted at neoblast 1, Spearman correlation with ordinal lineage order, and a label-shuffling compactness permutation test with 1,000 permutations and a maximum of 250 cells per group.

### Fixed-loading perturbation and enrichment

For each highly variable gene *g*, we estimated a fixed-loading perturbation in PCA space by adding *δℓ_g_* to each cell’s PC1–3 coordinate, where *ℓ_g_* denotes the loading vector of gene *g* on PC1–3. Because PCA was performed after gene-wise scaling, *δ* = 1.0 corresponds to a +1 standard deviation (SD) perturbation in standardized expression units propagated through the fixed PCA loading vector.

For each gene, perturbed PC1–3 coordinates were reprojected onto the unit sphere and compared with the original root-aligned spherical coordinates. Effects were summarized as progression-associated displacement,

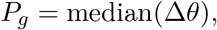

branch/state-position displacement,

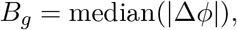

and radial displacement,

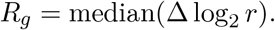

Here, positive *P_g_* indicates movement away from the root state, negative *P_g_* indicates movement toward the root state, *B_g_* captures lateral displacement around the root-aligned sphere, and *R_g_* captures changes in the pre-projection radial magnitude. These quantities represent computational displacements in fixed PCA-loading space and should not be interpreted as experimental perturbation effects.

Coordinate-specific gene classes were tested for pathway enrichment using Hallmark, Reactome, and Gene Ontology Biological Process libraries [38–42]. For Fig. 4, enrichment was computed using hypergeometric tests against the dataset-specific HVG background, with Benjamini–Hochberg false-discovery-rate correction implemented through Statsmodels [43]. For Fig. 3, enrichment was computed using gseapy.enrichr [44].

### Benchmarking and software

Benchmarking compared precomputed PC1–50 coordinates, PCA PC1–3, SPHERE-PCA-aligned PC1–3, 3D UMAP, 3D t-SNE, and a 3D scPhere latent embedding. scPhere was trained in normal latent mode for 10 epochs. Trustworthiness, continuity, and k-nearest-neighbor Jaccard overlap were computed with *k* = 15, a subsample size of *n* = 5,000, and seed sweep 0, 1, and 2, using scikit-learn routines where applicable [45]. Numerical analysis and plotting used NumPy, pandas, SciPy, Matplotlib, Statsmodels, Scanpy, and gseapy [36, 37, 43, 44, 46–48]. Package versions were: NumPy *≥* 2.0.2*, <* 3.0, pandas *≥* 2.3.3*, <* 3.0, SciPy *≥* 1.13.1*, <* 2.0, Matplotlib *≥* 3.9.4*, <* 4.0, Statsmodels *≥* 0.14.6*, <* 1.0, Scanpy *≥* 1.10.3*, <* 2.0, and gseapy *≥* 1.2.1*, <* 2.0. Enrichr libraries were accessed on May 11, 2026.

## Code availability and reproducibility

SPHERE-PCA is available as a Python package at https://github.com/imlong4real/SPHERE-PCA. Code for reproducing the manuscript figures is provided in the figure-generation module at https://github.com/imlong4real/SPHERE-PCA/tree/main/pca_sphere_projection/figures, with associated scripts, configuration files, and processed outputs included in the repository to support reproducibility.

## Acknowledgments

The authors thank Caroline Uhler, Alexis Battle, and Joshua Popp for their insights and feedback.

## Author Contributions

L.Y. and A.S.S. designed research; L.Y. performed research; L.Y., X.L., and M.L. analyzed data; A.D., S.C.H., J.M.T., and A.S.S. supervised the study; L.Y. wrote the paper; and all authors reviewed and approved the manuscript.

## Supplementary Information

## Supplementary Results

The supplementary analyses extend the main figures with stress tests, coordinate-gene annotation, targeted-panel analyses, and benchmarking. All analysis in this supplement follow the interpretive limits of the main text. SPHERE-PCA is treated as an interpretable geometric framework for dominant PC structure, not as a universal trajectory inference algorithm, a de novo lineage-discovery method, or a replacement for nonlinear embeddings.

### Planarian angular geometry

Supplementary Figure S1 compares angular relationships among planarian cell-type centroids in 3D SPHERE-PCA space with angular relationships computed from PC1-PC50 [24]. The 3D spherical representation provides a magnitude-free view of broad lineage-sector organization, while the PC50 angular matrix confirms that related global relationships are present in the higher-dimensional PC space.

### Coordinate gene classes

Supplementary Figure S2 supports the gene-coordinate analysis in Fig. 3 across human ESC differentiation, human germ cell, and human preimplantation embryo datasets [19, 20, 35]. Gene classes defined by association with root-aligned geodesic distance, pre-projection radial norm, or both were tested for gene-set enrichment using Gene Ontology Biological Process, Hallmark, and Reactome libraries [38, 39]. Overlap with the 97-gene Tirosh et al cell-cycle marker [25] list was evaluated by Fisher’s exact test against the HVG background with Benjamini-Hochberg FDR correction. These results support coordinate-specific biological consistency, not causal mechanism.

### Non-developmental and targeted spatial stress tests

Supplementary Figure S3 evaluates SPHERE-PCA in a breast cancer scRNA-seq atlas and targeted Xenium/TRACER spatial panels [30–33]. The breast cancer atlas shows cell-type-associated spherical regions. Xenium breast cancer and lung cancer panels show visible pre- and post-TRACER spherical structure, but the targeted panels contain approximately 300 genes. These analyses are stress tests for modality and panel design and are not interpreted as developmental trajectories.

### Embedding benchmark

Supplementary Figure S4 compares PCA (PC1-3), SPHERE-PCA, UMAP (3D), t-SNE (3D), and scPhere (3D) in the *C. elegans* embryogenesis atlas [1, 2, 14, 18]. Metrics include trustworthiness, continuity, k-nearest-neighbor Jaccard overlap, and absolute Spearman correlation between anchor distance and embryonic time. The benchmark supports complementary behavior: UMAP and t-SNE perform better on local-neighborhood metrics, while SPHERE-PCA provides an interpretable spherical coordinate system with competitive global developmental ordering.

**Figure S1:**
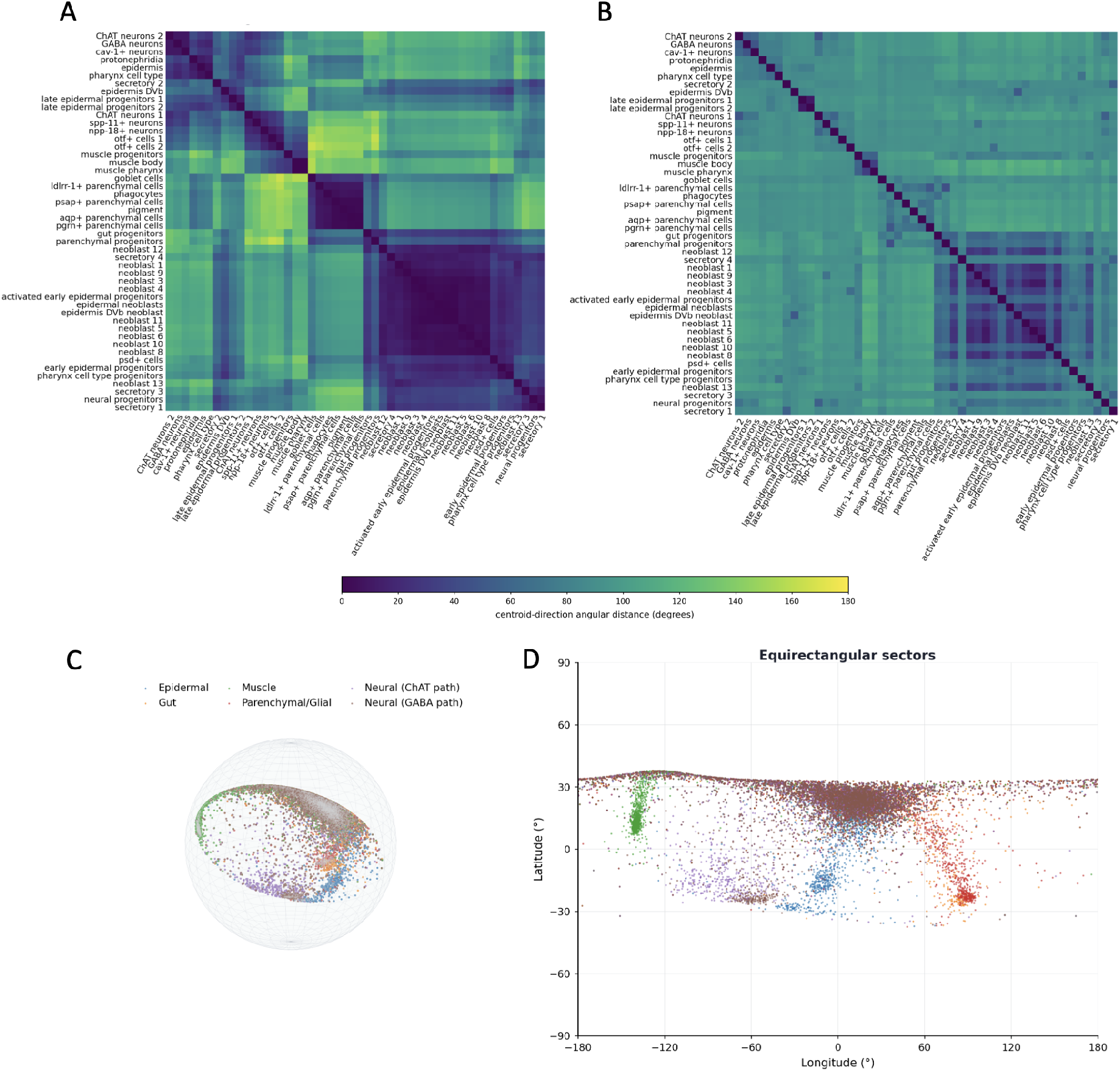
Planarian angular geometry preserves lineage-sector organization better than high-dimensional PC angular distances. (A) Heatmap of angular distances between planarian cell-type centroid directions computed from the 3D SPHERE-PCA representation. Each row and column corresponds to a cell type with at least the minimum cell count used by the pipeline; color indicates centroid angular distance. (B) Matched heatmap of angular distances between cell-type centroids computed in PC1-PC50 space using cosine distance converted to angular distance. (C) 3D SPHERE-PCA view colored by selected known planarian lineage groups, including epidermal, gut, muscle, parenchymal/glial, neural ChAT, and neural GABA paths, all rooted at neoblast 1. (D) Equirectangular projection of the same sphere and lineage coloring. The figure supports angular organization of published lineage annotations; it does not reconstruct or infer the planarian lineage tree.

**Figure S2:**
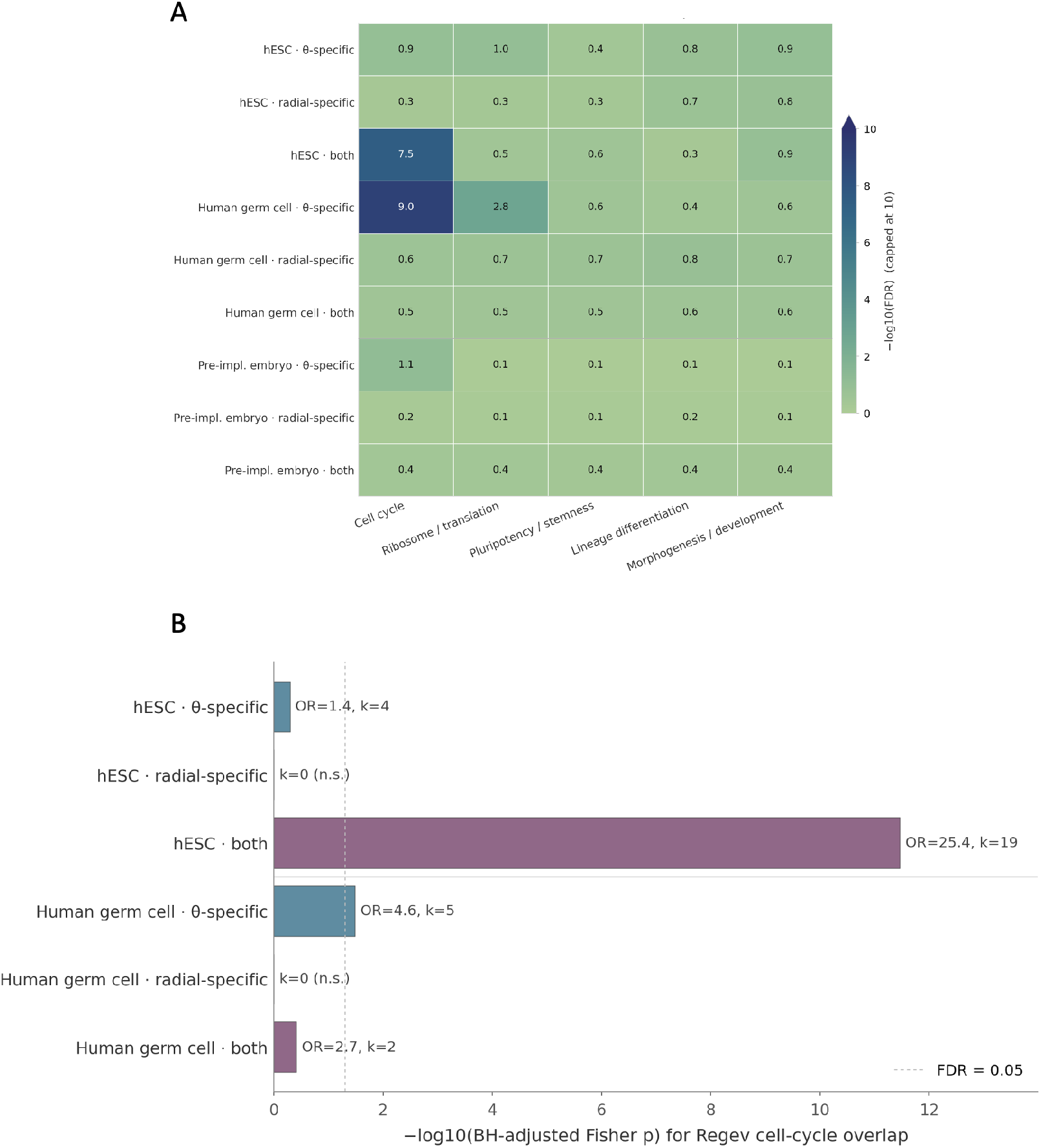
Coordinate-specific gene classes show pathway enrichment and overlap with canonical cell-cycle markers. (A) Heatmap of enrichment scores for *θ*-specific, radial-specific, and both-coordinate gene classes across selected biological categories in human ESC differentiation, human germ cell, and human preimplantation embryo datasets. Gene-set enrichment used Gene Ontology Biological Process, Hallmark, and Reactome libraries. Color indicates − log_10_ FDR, capped at 10, from hypergeometric or Enrichr-based over-representation analysis against the HVG background. (B) Overlap of coordinate-specific gene classes with the Regev/Tirosh 97-gene cell-cycle marker list. Bars show − log_10_ Benjamini-Hochberg-adjusted Fisher exact-test *P* values, with odds ratio and overlap count annotations where available. Strong overlap of hESC both-coordinate genes with cell-cycle markers supports activity-associated radial interpretation, but radial variation is not assumed to be cell cycle only.

**Figure S3:**
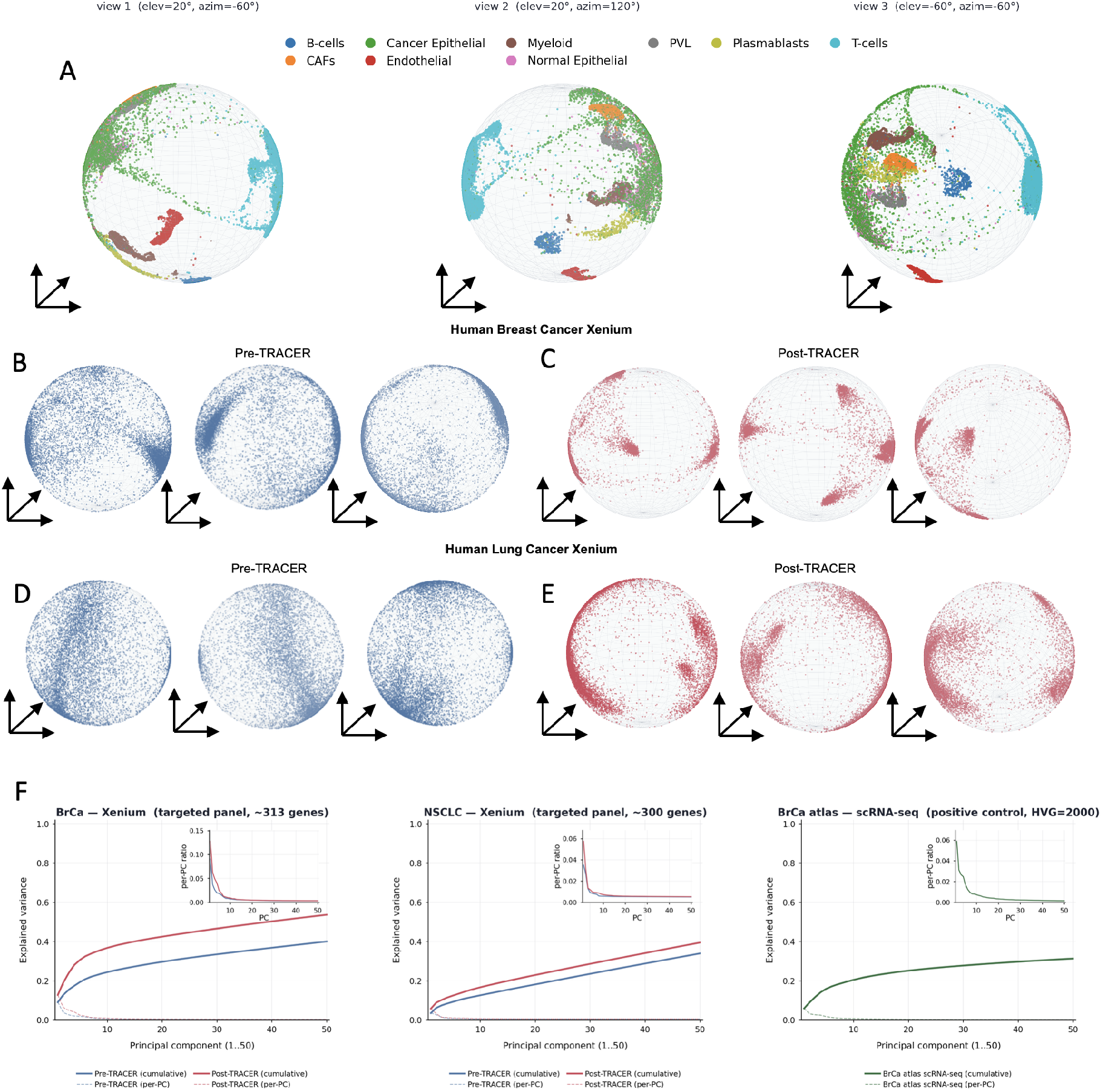
SPHERE-PCA structure in non-developmental and targeted spatial transcriptomic datasets. (A) Human breast cancer scRNA-seq atlas projected to the unit sphere and shown from three camera views. Cells are colored by major cell type; the observed regions reflect cell-type composition, not a developmental hierarchy. (B-E) Pre- and post-TRACER targeted Xenium breast cancer and lung cancer datasets projected to the unit sphere using all assayed genes and PC1-PC3, shown from multiple camera angles. Points are cells; colors follow the plotted cell-state or condition encodings in the generated figure tables. These targeted panels may show different PCA variance structure than whole-transcriptome data; spherical organization here reflects panel-dependent cell-state variation (F) PC1-PC50 explained-variance comparison for targeted Xenium panels and the breast cancer scRNA-seq atlas reference. Cumulative and per-PC variance curves show that targeted panels and whole-transcriptome scRNA-seq can have different PCA variance structure. This comparison quantifies differences in PCA variance structure between targeted and whole-transcriptome datasets and should not be used to calibrate interpretation of the Xenium embeddings above.

**Figure S4:**
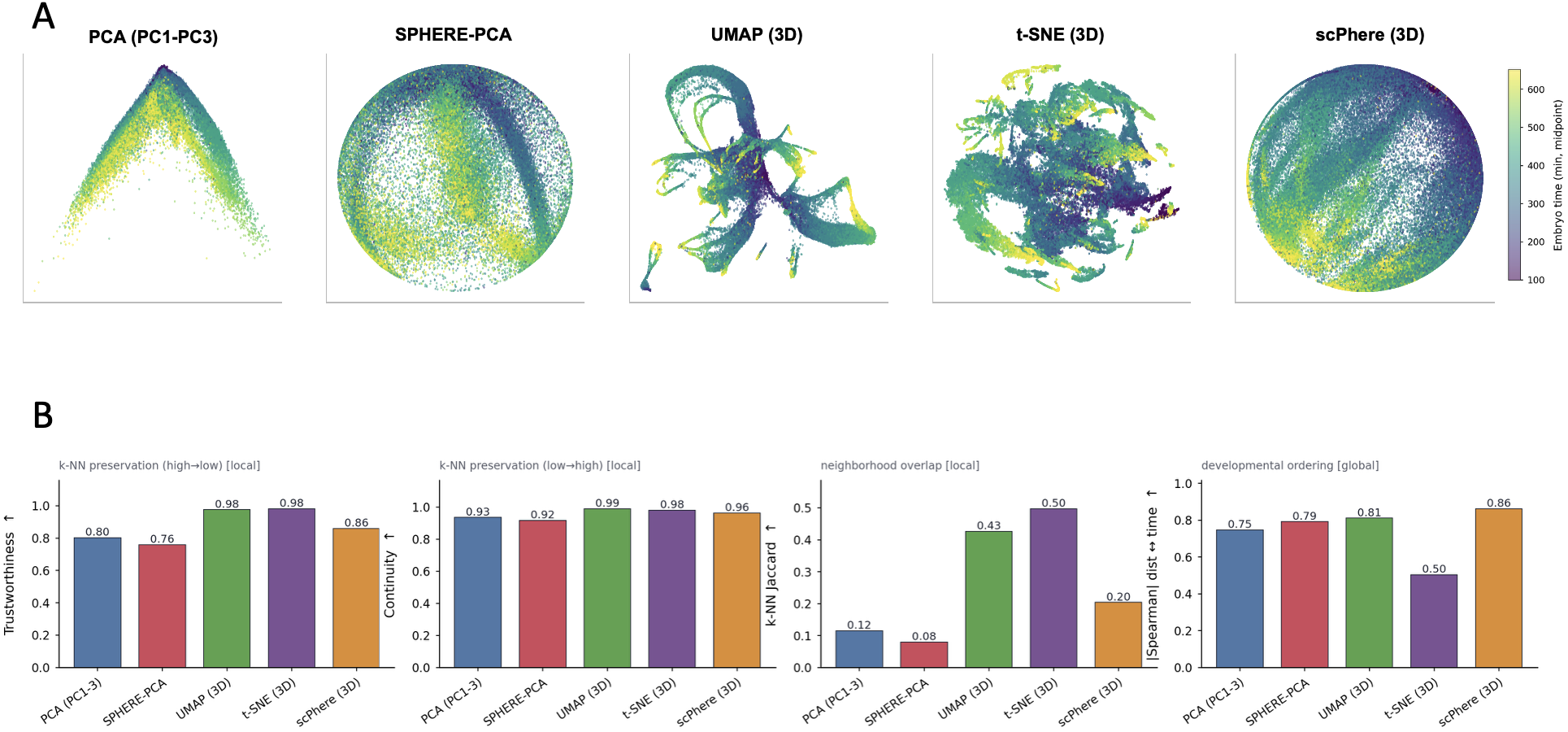
Benchmarking SPHERE-PCA against PCA, UMAP, t-SNE, and scPhere embeddings. (A) Visual comparison of 3D PCA, SPHERE-PCA, UMAP, t-SNE, and scPhere embeddings for the *C. elegans* embryogenesis atlas, with cells colored by embryonic time. (B) Quantitative benchmark metrics against a PC1-PC50 reference: trustworthiness, continuity, k-nearest-neighbor Jaccard overlap, and absolute Spearman correlation between anchor distance and embryonic time. SPHERE-PCA provides an interpretable spherical coordinate system and competitive global developmental ordering, while UMAP and t-SNE show higher trustworthiness and continuity, consistent with their local-neighborhood objectives; scPhere shows higher Spearman correlation between anchor distance and embryonic time.

**Movie S1. Rotating SPHERE-PCA visualization of the *C. elegans* embryogenesis atlas.** Cells from the *C. elegans* embryogenesis atlas are shown on the unit sphere after PC1-PC3 *L*_2_ projection and root alignment using the 100-130 embryonic time bin. Color indicates embryonic time bin. The movie is a visualization aid for the spherical geometry and should not be interpreted as an independent embedding or an automated lineage-detection analysis.

**Movie S2. Rotating SPHERE-PCA visualization of the planarian atlas.** Planarian cells are shown on the unit sphere after SPHERE-PCA projection and alignment to the neoblast reference used in the planaria analyses. Color follows the atlas or lineage-state encoding in the generated movie. The movie supports visual inspection of angular organization for known planarian annotations but does not claim de novo lineage reconstruction.

